# Structure and functional mapping of the KRAB-KAP1 repressor complex

**DOI:** 10.1101/2022.03.17.484746

**Authors:** Guido A. Stoll, Ninoslav Pandiloski, Christopher H. Douse, Yorgo Modis

**Author notes:** Correspondence (Y.M.).

## Abstract

Transposable elements (TEs) are a genetic reservoir from which new genes and regulatory elements can emerge. Expression of TEs can be pathogenic, however, and is tightly regulated. KRAB domain-containing zinc finger proteins (KRAB-ZFPs) recruit co-repressor KRAB-associated protein 1 (KAP1/TRIM28) to regulate many TEs but how KRAB-ZFPs and KAP1 interact remains unclear. Here, we report the crystal structure of the KAP1 tripartite motif (TRIM) in complex with the KRAB domain from a human KRAB-ZFP, ZNF93. Structure-guided mutations in the KAP1-KRAB binding interface abolished repressive activity in an epigenetic TE silencing assay. Deposition of H3K9me3 over thousands of loci is lost genome-wide in cells expressing a KAP1 variant with mutations that abolish KRAB binding. Our work identifies and functionally validates the KRAB-KAP1 molecular interface, which forms the nexus of a transcriptional control axis vital to vertebrates and underpins programmable gene repression by CRISPRi.

## Introduction

More than half of the human genome consists of transposable elements (TEs) (Friedli & Trono, 2015). TEs can be acquired when viral DNA integrates into the genome of a host germline cell. These endogenous viral elements (EVEs) can retain the ability to replicate by expressing the viral reverse transcriptase and integrase, which convert EVE transcripts into DNA and integrate the DNA into the host genome (Friedli & Trono, 2015). Other TEs, such as LINEs (long interspersed nuclear elements), replicate via a similar retrotransposition mechanism, but the distinct sequence and biochemical activities of LINE proteins suggest they evolved from early eukaryotic, rather than viral genetic elements (Goodier, 2016). Approximately 100 LINEs are replication-competent and 2-5% of newborn children have a new LINE insertion (Goodier, 2016). Although most human EVEs have lost transposition activity, some of the most recently acquired human endogenous retrovirus (HERVs) retain the potential to be transcribed, translated, and transposed (Grow *et al*, 2015; Li *et al*, 2015).

Transcription, translation, and transposition of TEs are potentially pathogenic, particularly in embryogenesis, chronic infection, and stress responses, when pro-transcriptional chromatin modifications are enriched (Azebi *et al*, 2019). Accumulation of TE-derived nucleic acids is associated with autoimmune diseases including geographic atrophy, lupus and Sjögren’s syndrome (Goodier, 2016; Hung *et al*, 2015). Aberrant expression of HERV proteins is associated with cancer and neurodegeneration (Kremer *et al*, 2019; Li *et al*., 2015). When transposition events disrupt tumor suppressor genes or enhance oncogene expression they contribute to cancer (Hancks & Kazazian, 2016; Lamprecht *et al*, 2010). Gene disruption by TE transposition is also linked to genetic disorders such as hemophilia and cystic fibrosis (Hancks & Kazazian, 2016). The activities of TEs must therefore be tightly regulated.

Counterbalancing these risks, TEs are a genetic reservoir from which new genes and regulatory elements can emerge. TEs drive the evolution of transcriptional networks by spreading transcription factor binding sites, promoters and other regulatory elements (Chuong *et al*, 2016; Friedli & Trono, 2015). Pluripotency-associated transcription factors required for cell fate determination bind to sites within TEs (Friedli & Trono, 2015). Some TE genes have also been coopted to fulfill important cellular functions. For example, TE-derived proteins catalyze V(D)J recombination (Zhou *et al*, 2004), trophoblast fusion in placental development (Dupressoir *et al*, 2012; Friedli & Trono, 2015), and cell-to-cell mRNA transfer required for synaptic plasticity and memory formation (Pastuzyn *et al*, 2018).

The primary mechanism cells have evolved to control TEs is epigenetic transcriptional silencing. In tetrapod vertebrates, Krüppel-associated box zinc-finger proteins (KRAB-ZFPs) and KRAB-associated protein 1 (KAP1, also known as TRIM28 or TIF1β), are key repressors of TEs (Helleboid *et al*, 2019; Rowe *et al*, 2010). KRAB-ZFPs are the largest mammalian transcription factor family. Humans have 350-400 KRAB-ZFPs, the majority of which recognize specific TE-derived DNA sequences with their tandem variable zinc-finger arrays (Imbeault *et al*, 2017; Jacobs *et al*, 2014). The expansion in the number of KRAB-ZFPs in mammals has been attributed to evolutionary pressure from TEs mutating to escape recognition, resulting in an arms race between hosts and TEs (Jacobs *et al*., 2014). Because many sequences targeted by KRAB-ZFPs have been repurposed as promoters or enhancers during the course of evolution, some of the older KRAB-ZFPs now regulate physiologically important processes such as genomic imprinting, embryogenesis, brain development, and immunity (Azebi *et al*., 2019; Imbeault *et al*., 2017; Johansson *et al*, 2022; Li *et al*, 2021; Tie *et al*, 2018; Tycko *et al*, 2020). Once bound to DNA, (all but a few) KRAB-ZFPs recruit corepressor KAP1 via the conserved KRAB domain (Friedman *et al*, 1996; Helleboid *et al*., 2019; Kim *et al*, 1996; Moosmann *et al*, 1996; Tycko *et al*., 2020). Disrupting KAP1 function is lethal early in embryonic development (Cammas *et al*, 2000).

KRAB domains typically contain a KRAB-A box (40-50 amino acids) necessary and sufficient for KAP1-dependent repression, and a KRAB-B box (20-25 amino acids) with an accessory role (Margolin *et al*, 1994; Peng *et al*, 2007; Tycko *et al*., 2020; Witzgall *et al*, 1994). Solution NMR studies of the KRAB-A box from a mouse KRAB-ZFP generated a partly α-helical, partly disordered structural ensemble (Saito *et al*, 2003). A complete mutational scan of the KRAB domain identified 12 residues in the KRAB-A box where mutations abolished silencing, along with a few residues where substitutions enhanced silencing (Tycko *et al*., 2020). Most of the residues required for silencing were required for KAP1 binding in a recombinant protein binding assay (Peng *et al*, 2009) and predicted to cluster together in a structural model of the KRAB-A box (Tycko *et al*., 2020). KRAB domains with the strongest KAP1 binding and silencing activities are among the most powerful transcriptional repressors. Fused to inactive Cas9 in the CRISPR interference (CRISPRi) approach, KRAB domains allow potent programmable gene repression (Alerasool *et al*, 2020; Gilbert *et al*, 2014; Thakore *et al*, 2015).

At its target DNA loci, KAP1 functions as a recruitment platform for repressive chromatin-modifying enzymes including histone H3K9 methyltransferase SETDB1, heterochromatin protein HP1, and the nucleosome remodeling and deacetylase (NuRD) complex (Schultz *et al*, 2002; Schultz *et al*, 2001). We showed previously that KAP1 dimerizes via the coiled-coil domain in its RING, B-box zinc finger and Coiled-Coil (RBCC) motif (Stoll *et al*, 2019). One KAP1 dimer binds a single KRAB domain. A set of four structure-based mutations in the coiled-coil domain, near the dyad of the KAP1 RBCC dimer, abolished KRAB binding and transcriptional silencing, suggesting these mutations map to the KRAB binding site (Stoll *et al*., 2019). However, the structure of the KRAB-KAP1 interaction remains unknown. Here, we report the crystal structure of the KAP1 RBCC in complex with the KRAB domain from ZNF93, a KRAB-ZFP that represses LINE-1 elements in primates (Jacobs *et al*., 2014). The structure provides a three-dimensional atlas of the KRAB-KAP1 binding interface. We use an epigenetic gene silencing assay to confirm that KAP1 residues forming key contacts with the KRAB domain are essential for silencing. Epigenomic profiling shows that deposition of repressive H3K9me3 marks is affected genome-wide in cells expressing a KAP1 variant with mutations at the KRAB binding interface. Our work identifies and functionally validates the KRAB-KAP1 molecular interface, at the nexus of a transcriptional control axis that is vital to vertebrates and underpins programmable gene repression by CRISPRi.

## Results

### Crystallographic structure determination of a KRAB-KAP1 core complex

To elucidate the molecular basis of transcriptional regulation by KRAB-ZFPs, we determined the crystal structure of the KRAB domain from human ZNF93, a KRAB-ZFP that represses LINE-1 elements (Jacobs *et al*., 2014), in complex with the RBCC domain of KAP1. Structure determination was technically challenging. Initial crystallization trials with the ZNF93 KRAB–KAP1 RBCC complex were unsuccessful. Fusion of bacteriophage T4 lysozyme to the N-terminus of KAP1 RBCC induced crystallization, but the crystals diffracted X-rays poorly, precluding structure determination. The crystal quality was improved by deleting the flexible B-box 1 domain of KAP1 and adding the CUE1 domain from the chromatin remodeler SMARCAD1, which binds to the coiled-coil domain of KAP1 (Ding *et al*, 2018; Lim *et al*, 2019), to the complex (**Fig 1A**). The resulting crystals allowed collection of X-ray diffraction data extending to 2.8 Å resolution (**Table 1**). The structure was determined by molecular replacement (see **Materials and Methods**), but the electron density in the KRAB domain, particularly for side chains, was weaker than for the rest of the complex (**Fig 1B**). The directionality and sequence register of the KRAB backbone remained too ambiguous to allow an atomic model to be built *de novo*. However, guided by the AlphaFold2 model of the ZNF93 KRAB domain alone and the NMR structure of the KRAB-A box from a mouse KRAB-ZFP (UniProt A0A087WRJ1; (Saito *et al*., 2003)), we were able to build an initial atomic model of the ZNF93 KRAB-A box bound to KAP1 (**Fig 1C-D**). To validate this model, we introduced methionine point mutations throughout the KRAB domain, generating the ZNF93 variants I11M, C20M, L28M and L40M. The corresponding residues in the ZNF10 KRAB domain were previously shown to tolerate mutation to methionine without loss of silencing function (Tycko *et al*., 2020). Selenomethionine derivatives of these four ZNF93 mutants were purified in complex with KAP1 RBCC and crystallized. X-ray diffraction data were collected (at the selenium K absorption edge) and anomalous Fourier maps calculated. For each mutant, the position of the newly introduced selenium site was successfully located in the anomalous maps, allowing unambiguous identification of the mutated residue and confident sequence assignment (**Figs 1E and S1**).

**Figure 1.**
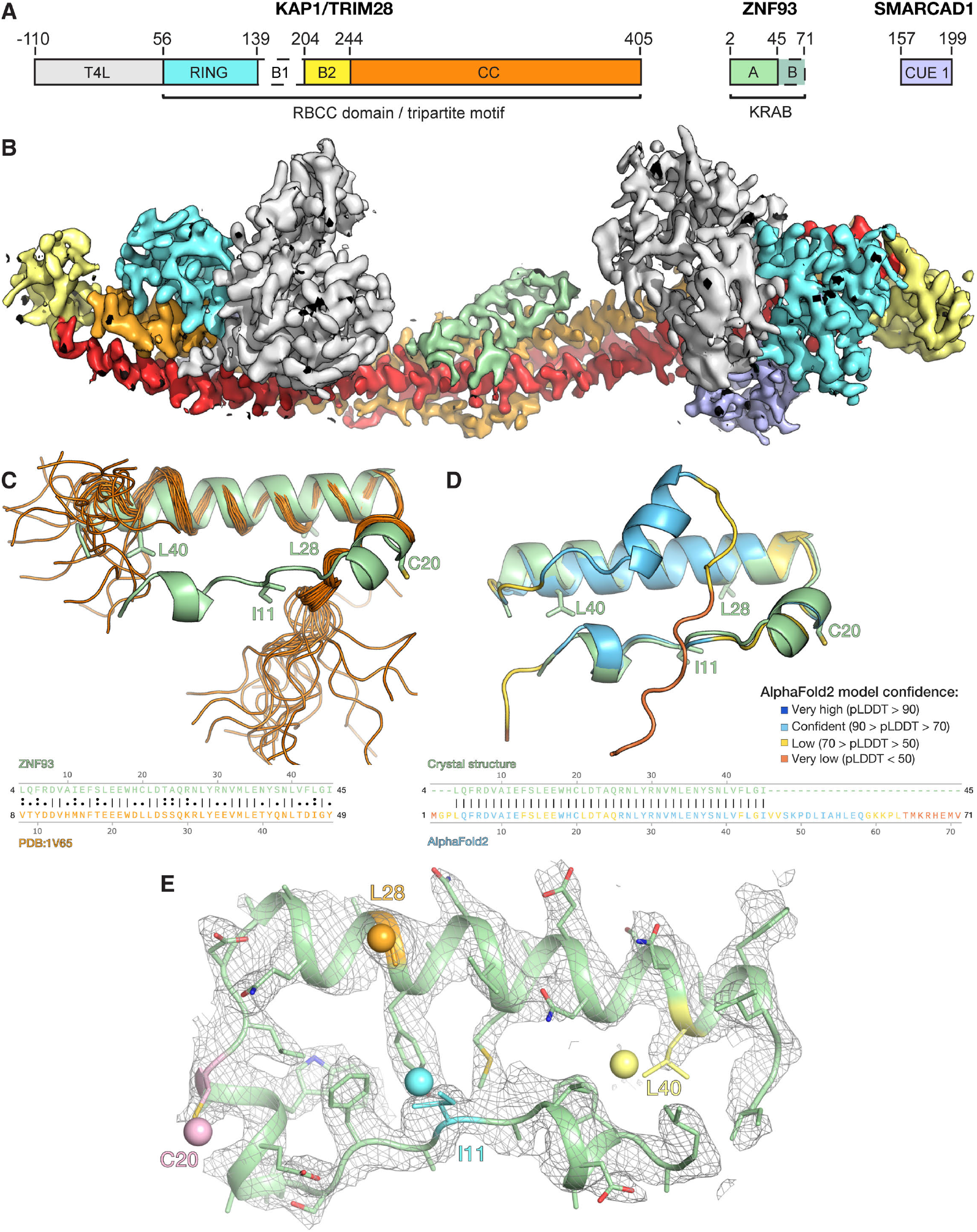
Structure determination of the ZNF93-KAP1 core complex. See also **Fig S1**. **(A)** Domain organisation of the crystallized complex. T4L, T4 lysozyme; B1, B-box 1; B2, B-box 2; CC, coiled-coil. **(B)** 2*F*_o_ - *F*_c_ electron density map for one asymmetric unit. The map is contoured at 1 σ and colored by domain as in (*A*), except the second CC, which is in red. **(C)** Superposition of the crystal structure of ZNF93 KRAB-A (green) on the solution NMR structure of the KRAB-A box from a mouse KRAB-ZFP (UniProt A0A087WRJ1; orange; (Saito *et al*., 2003)). A sequence alignment of the two domains is shown below. **(D)** Superposition of the ZNF93 KRAB-A crystal structure (green) and the AlphaFold2 prediction (Jumper *et al*, 2021; Varadi *et al*, 2022) of ZNF93 KRAB (colored by confidence score, pLDDT). A sequence alignment of the two domains is shown below. **(E)** Close-up of the electron density (2*F*_o_ - *F*_c_ map) for the wild-type ZNF93 KRAB-A domain, contoured at 1.0 σ. Residues chosen for mutation to methionine are highlighted in cyan (I11), pink (C20), orange (L28) or yellow (L40). The selenium sites located for each of these variants in anomalous Fourier maps are shown as spheres.

**Table 1.** Crystallographic data collection, refinement and structure validation parameters and statistics for complexes containing ZNF93 KRAB, T4 lysozyme fused KAP1 RBCC(ΔB1), and SMARCAD1 CUE1. See also **Figs 1 and S1**.

|  | Wild-type<br>Native | ZNF93 I11M<br>SeMet | ZNF93 C20M<br>SeMet | ZNF93 L28M<br>SeMet | ZNF93 L40M<br>SeMet |
| --- | --- | --- | --- | --- | --- |
| **Data collection** |  |  |  |  |  |
| Space group | *C* 1 2 1 | *C* 1 2 1 | *C* 1 2 1 | *C* 1 2 1 | *C* 1 2 1 |
| Cell dimensions<br>a, b, c (Å)<br>$\alpha$, $\beta$, $\gamma$ (°) | 190.3, 68.6, 149.1<br>90, 114.0, 90 | 186.8, 69.9, 156.1<br>90, 113.6, 90 | 187.2, 69.8, 156.9<br>90, 113.8, 90 | 187.3, 70.0, 156.8<br>90, 113.8, 90 | 187.9, 69.8, 157.7<br>90, 113.9, 90 |
| Wavelength (Å) | 0.9795 | 0.97949 | 0.9795 | 0.97951 | 0.9795 |
| Resolution (Å)<sup>‡</sup> | 61 - 2.8 (2.9 - 2.8) | 52 - 3.5 (3.63 - 3.5) | 52 - 3.3 (3.42 - 3.3) | 51 - 3.5 (3.63 - 3.5) | 63 - 3.5 (3.63 - 3.5) |
| Observations<sup>‡</sup> | 119,389 (12,179) | 91,722 (8,800) | 10,8241 (10,531) | 91,049 (8,870) | 90,349 (8,968) |
| Unique reflections<sup>‡</sup> | 42,537 (4,206) | 23,605 (2,323) | 28,280 (2,786) | 23,798 (2,324) | 23,851 (2,347) |
| $R_{merge}$<sup>‡</sup> | 0.0747 (0.50) | 0.129 (0.67) | 0.126 (1.0) | 0.157 (1.0) | 0.115 (0.73) |
| $R_{pim}$<sup>‡</sup> | 0.0518 (0.34) | 0.0751 (0.39) | 0.0743 (0.61) | 0.0917 (0.60) | 0.0681 (0.43) |
| $\langle I \rangle / \sigma I$<sup>‡</sup> | 21 (2.9) | 13 (1.8) | 14 (1.8) | 12 (1.8) | 9 (1.9) |
| Completeness (%)<sup>‡</sup> | 97.3 (97.3) | 99.2 (99.0) | 99.5 (98.9) | 99.3 (98.7) | 99.2 (99.5) |
| Multiplicity<sup>‡</sup> | 2.8 (2.9) | 3.9 (3.8) | 3.8 (3.8) | 3.8 (3.8) | 3.8 (3.8) |
| CC(1/2)<sup>‡</sup> | 0.99 (0.70) | 0.99 (0.87) | 0.99 (0.58) | 0.97 (0.68) | 0.98 (0.87) |
| **Refinement** |  |  |  |  |  |
| $R_{work} / R_{free}$ | 0.2255 / 0.2728 | 0.2288 / 0.2696 | 0.2547 / 0.3083 | 0.2571 / 0.3051 | 0.2430 / 0.2941 |
| N. of non-H atoms |  |  |  |  |  |
| Protein | 7,889 | 8,167 | 8,133 | 8,163 | 8,213 |
| Zn<sup>2+</sup> Ions | 8 | 8 | 8 | 8 | 8 |
| Solvent | 0 | 0 | 0 | 0 | 0 |
| Mean B-factor (Å<sup>2</sup>) | 69 | 141 | 135 | 156 | 152 |
| Clashscore* | 8.98 | 2.44 | 2.45 | 3.11 | 3.40 |
| RMS deviations |  |  |  |  |  |
| Bond lengths (Å) | 0.013 | 0.003 | 0.003 | 0.003 | 0.003 |
| Bond angles (°) | 1.6 | 0.68 | 0.71 | 0.85 | 0.73 |
| Ramachandran plot |  |  |  |  |  |
| % favored | 97.2 | 96.5 | 96.6 | 95.8 | 96.5 |
| % allowed | 2.3 | 2.9 | 3.0 | 3.7 | 2.8 |
| % outliers | 0.5 | 0.6 | 0.4 | 0.5 | 0.7 |
| PDB code | 7Z36 |  |  |  |  |

<sup>‡</sup>Numbers in parentheses refer to the highest resolution shell.
\*Clashscore was calculated with Phenix v1.20 (Adams *et al*, 2011).

### Structure of a KRAB domain in complex with KAP1

The structure contains one KAP1 RBCC dimer, two SMARCAD1 CUE1 domains and one ZNF93 KRAB domain in the crystallographic asymmetric unit (**Fig 2**). Consistent with previous biochemical studies (Stoll *et al*., 2019), a single KRAB domain binds the KAP1 RBCC dimer, near the twofold axis, in the central region of the coiled-coil domain. Contacts with KAP1 are exclusively mediated by the KRAB-A box (residues 3-44 of ZNF93), while the KRAB-B box (residues 45-71) is disordered and not visible in the electron density map. This is consistent with previous observations that the KRAB-A box is sufficient for KAP1 binding and transcriptional repression, whereas KRAB-B has an accessory function and is absent in some KRAB-ZFPs (Margolin *et al*., 1994; Peng *et al*., 2007; Tycko *et al*., 2020; Witzgall *et al*., 1994). The KRAB-A box backbone is U-shaped and contains three α-helical segments: a single helical turn at the N-terminus (α1, residues 6-9), a short central helix (α2, residues 15-20) and a longer C-terminal helix (α3, residues 23-43). The overall conformation of the KRAB-A box in the refined crystal structure remains similar to the AlphaFold2 model of the ZNF93 KRAB and the KRAB-A box NMR structure that guided model building (**Fig 1A-B**).

**Figure 2.**
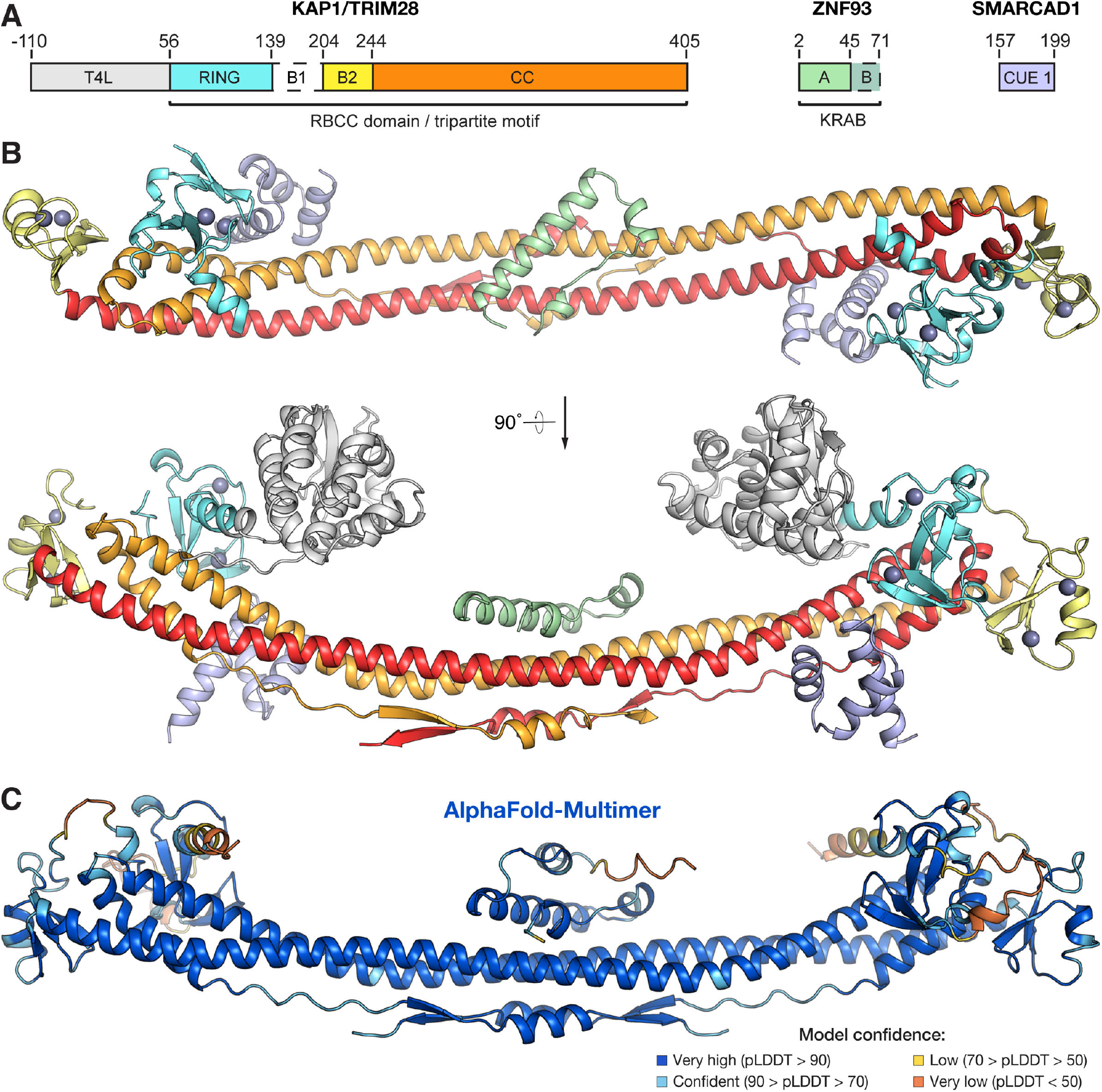
Structure of the ZNF93 KRAB domain of bound to KAP1 RBCC. **(A)** Domain organisation of the crystallized complex (as in **Fig 1A**). **(B)** Overall structure of the KRAB-KAP1 complex. Domains are coloured as in *(A)*. Zn atoms are shown as grey spheres. **(C)** Structure of ZNF93 KRAB:KAP1 RBCCΔB1 complex predicted by AlphaFold-Multimer (Evans *et al*., 2021; Mirdita *et al*., 2021)).

The AlphaFold2 model spans the entire KRAB domain, including a two-turn helix and random coil for the KRAB-B box, but the confidence score for most of the KRAB-B box is low (pLDDT < 70). In the NMR structure, the N- and C-termini of the KRAB-A box and the entire KRAB-B box are unstructured and highly flexible, consistent with biophysical data showing that KRAB domains are largely unstructured in isolation (Peng *et al*., 2007). We conclude that KRAB domains are mostly disordered prior to binding KAP1 (with some α-helical character in the KRAB-A box), and that the N- and C-termini of the KRAB-A box adopt a conserved, largely α-helical fold upon binding KAP1.

The RBCC domain of KAP1 in contrast displays no significant conformational changes in response to KRAB binding. The RING, B-box 2, and bound SMARCAD CUE1 domains are all distal from the KRAB-KAP1 interaction interface and unaffected by KRAB binding. The KAP1 coiled-coil domain has a higher curvature in the KRAB-KAP1 complex structure than in previously determined crystal structures of the KAP1 RBCC domain. However, the curvature of the coiled-coil domain varied in different crystal structures of KAP1 RBCC, demonstrating a degree of flexibility. Hence, it is unclear to what extent the crystal packing or KRAB binding contribute to the increased coiled-coil curvature.

After we completed and refined the crystal structure, AlphaFold-Multimer became available (Evans *et al*, 2021; Mirdita *et al*, 2021). We used it to predict the structure of a 1:2 ZNF93 KRAB:KAP1 RBCC complex. The resulting model was remarkably similar to our crystal structure (**Fig 2C**; Rmsd 3.1 Å), providing mutual validation of the two models.

### Key interacting residues in KRABs and KAP1 are highly conserved

KRAB domains bind to KAP1 with high affinity, with dissociation constants in the low nanomolar range (Stoll *et al*., 2019). Our structure of ZNF93 KRAB bound to KAP1 reveals that the interaction is mostly hydrophobic in nature, with a buried surface area of 1,108 Å^2^. The KAP1-binding surface of the KRAB domain is enriched in exposed hydrophobic amino acids and recognizes a hydrophobic patch in the central region of the KAP1 coiled-coil domain (**Fig 3A**). The hydrophobic core of the KRAB-KAP1 interface is surrounded by a network of polar interactions and salt bridges (**Fig 3A**). The KRAB forms contacts with both subunits of the KAP1 dimer. Hence, despite having an essentially identical structure, each KAP1 subunit binds to different regions of the KRAB (**Fig 3**).

**Figure 3.**
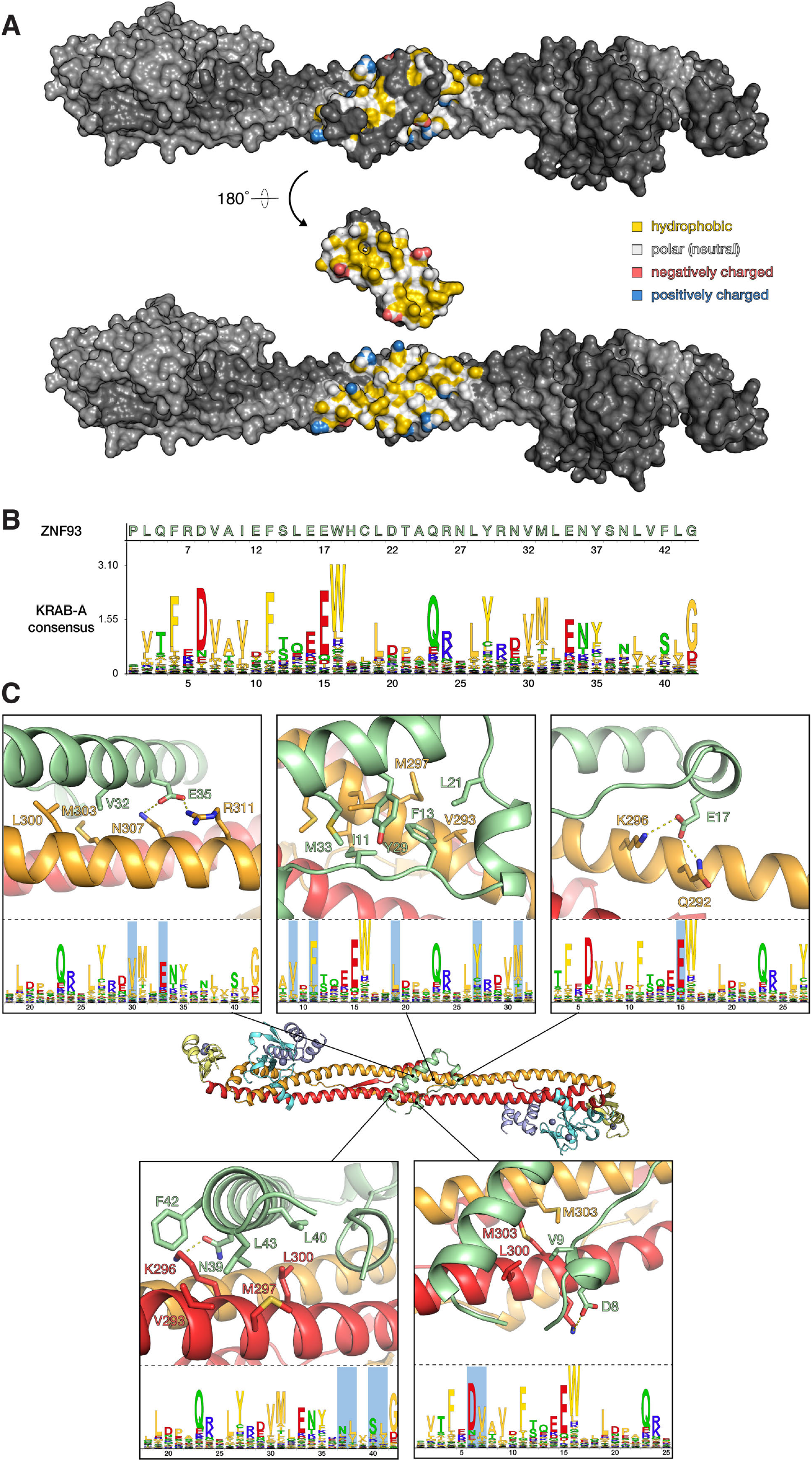
Molecular details of the KRAB-KAP1 interface. See also **Fig S2**. **(A)** The KRAB-KAP1 complex is shown in surface representation, with the KRAB domain in its natural orientation (*top*) or rotated by 180° to reveal the interaction surface contacting KAP1 (*bottom*). Residues in the interface are coloured according to their atomic properties using the YRB scheme (Hagemans *et al*, 2015). **(B)** KRAB domain Hidden Markov Model (HMM) logo (Mistry *et al*, 2021). **(C)** Close-up views of key residues in the KRAB-KAP1 interface. The corresponding positions in the KRAB consensus sequence are highlighted in the lower panels.

The amino acid sequences of KRAB-ZFP KRAB-A boxes are highly conserved across tetrapod vertebrates, except for a small KRAB-ZFP subset, which is thought to have acquired KAP1-independent functions (Helleboid *et al*., 2019). Fused to catalytically inactive Cas9, the KRAB domains of ZNF10 and ZIM3 allow potent gene repression that can be programmed in a sequence-specific manner via the CRISPRi approach (Alerasool *et al*., 2020; Gilbert *et al*., 2014; Margolin *et al*., 1994; Thakore *et al*., 2015). Deep mutagenesis studies of ZNF10 have identified the residues essential for KRAB repressor function (Tycko *et al*., 2020). These residues are generally conserved in the KRAB-ZFP family and across species. Our structure of the KRAB-KAP1 complex provides a mechanistic explanation for these observations. For example, mutation of the highly conserved Asp8 and Val9 residues in the KRAB-A box results in loss of silencing. Both residues form direct contacts with KAP1 in the KRAB-KAP1 structure, with Val9 buried in the hydrophobic core and Asp8 forming a salt bridge with Arg304 of KAP1 (**Fig 3B**). As noted above, hydrophobic contacts form the core of the KRAB-KAP1 interface. Besides Val9, these hydrophobic contacts involve Ile11, Phe13, Leu21, Tyr29, Val32 and Met33 in the KRAB domain, all of which are conserved and required for silencing (Tycko *et al*., 2020). Phe42 forms weak hydrophobic contacts with several neighboring residues, both in KAP1 and within the KRAB. The ZNF10 KRAB, used in first-generation CRISPRi, has a serine in the equivalent position (Ser51), and mutation of this serine to phenylalanine increases repression activity (Tycko *et al*., 2020). Notably, the KRAB domain with the highest reported repression potency in CRISPRi, the ZIM3 KRAB (Alerasool *et al*., 2020), also has a serine at this position (Ser46; **Fig S2**), suggesting that the potency of ZIM3 could be further increased by the substitution S46F.

Among the residues forming the ring of polar interface contacts, mutation of Glu17, one of the most conserved residues in the KRAB domain, to any other amino acid results abolishes KAP1 silencing (Tycko *et al*., 2020). This key KRAB domain residue forms a salt bridge with Lys296 and a hydrogen bond with Gln292 of KAP1 (**Fig 3B**). Similarly, Glu35, which is also essential for silencing, forms contacts with two residues in the KAP1 coiled-coil domain: a salt bridge with Arg311 and a hydrogen bond with Asn307. To assess the contribution of this salt bridge to KRAB1-KAP1 complex formation, we generated the charge reversal mutants ZNF93 E35R and KAP1 R311E and measured the binding constants of resulting KRAB-KAP1 complexes by surface plasmon resonance (SPR). The mutation E35R in ZNF93 KRAB reduced its binding affinity for wild-type KAP1 26-fold (from a *K*_d_ of 7.5 nM to 198 nM; **Fig 4A**). Combining the ZNF93 KRAB E35R and KAP1 R311E variants, which restored charge complementarity, partially restored binding affinity (**Fig 4A**).

**Figure 4.**
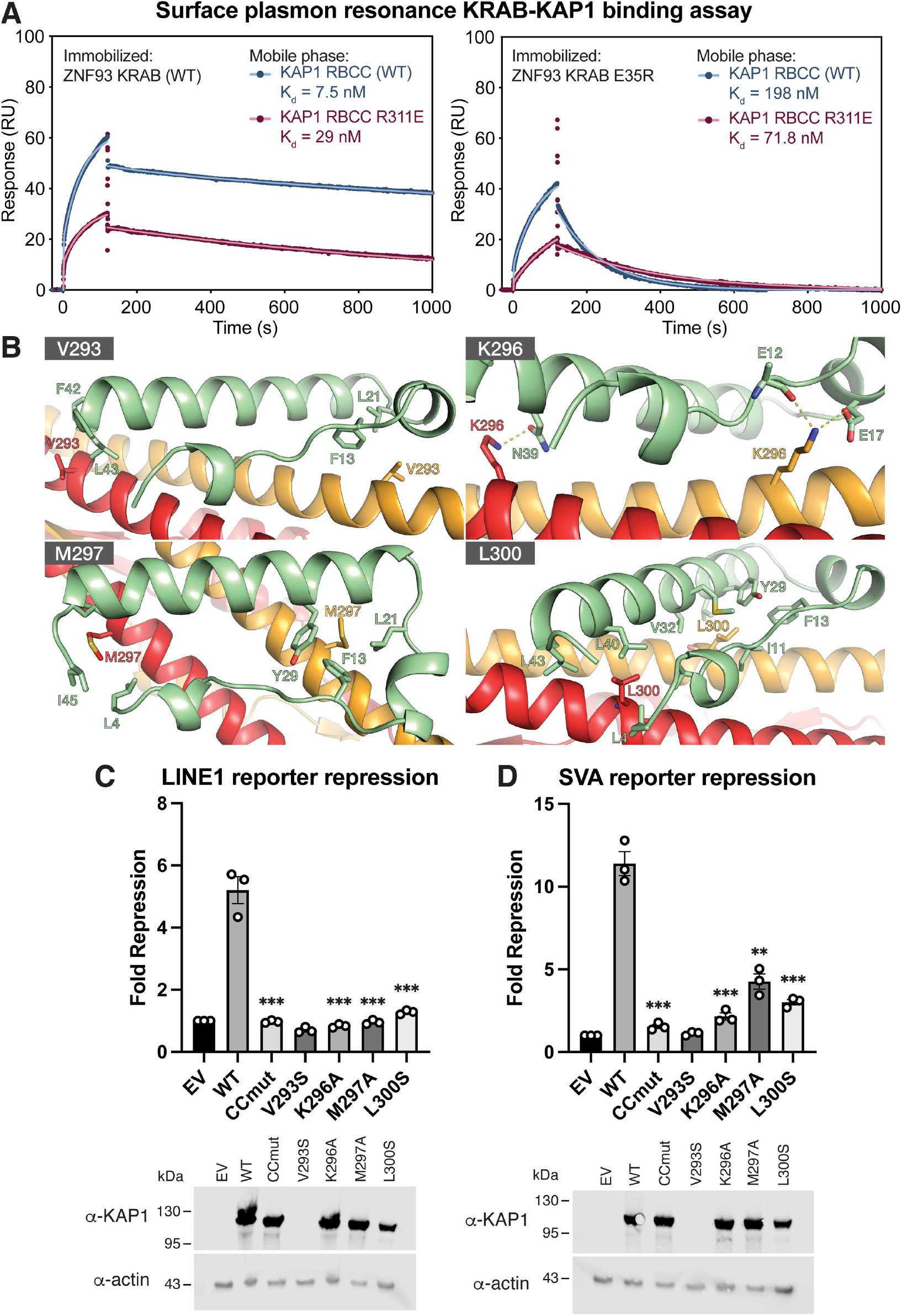
Effects of point mutations in the KRAB binding site and KAP1 binding and silencing. See also **Fig S3**. **(A)** Surface plasmon resonance (SPR) KAP1-KRAB binding assay. MBP-KRAB was immobilized on the chip. WT or R311E KAP1 RBCC were flowed over the chip. Data points are shown in dark red or blue; fits are shown as light red or blue lines. See **Fig S3** for binding kinetics constants. **(B)** Position of the mutations in the KRAB-KAP1 interface. **(C)** LINE-1 reporter repression with single point mutants and a KAP1 variant combining all four of these mutations (CCmut; V293S/K296A/M297A/L300S) **(D)** SVA reporter repression with the same set of mutants as in *(B)*. Data were normalized to KAP1 KO cells transfected with an empty vector (EV). Data are presented as fold-repression of reporter luciferase luminescence. Error bars represent standard error of the mean between measurements (n = 3). Statistical significance was assigned as follows: not significant (ns), P > 0.05; *, P < 0.05; **, P < 0.01; ***, P < 0.001. Lower panels: Western blots of cell lysates from KAP1 KO HEK293T cells transfected with each of the variants or empty vector.

The C-terminal KRAB-A sequence is more variable, but a strong preference for aliphatic residues at positions 40 and 43 is explained by the structure as these residues form part of the hydrophobic core of the KRAB-KAP1 interface. Charged side chains at these positions disrupt silencing (Tycko *et al*., 2020).

### Mutations in the KRAB-KAP1 interface abolish repression of L1 and SVA reporters

Our KRAB-KAP1 structure identifies the complete set of residues that form contacts at the binding interface. To assess the importance of individual key contacts, we measured the effects of structure-based interface mutations on KRAB-dependent transcriptional silencing in a dual luciferase reporter assay. We used previously described reporter constructs in which sequences from LINE-1 element repressed by ZNF93 or an SVA-D element repressed by ZNF91 were cloned upstream of firefly luciferase (Jacobs *et al*., 2014; Robbez-Masson *et al*, 2018). KAP1-knockout (KO) HEK 293T cells were cotransfected with the reporter plasmid and plasmids encoding ZNF93 or ZNF91, WT or mutant KAP1, and *Renilla* luciferase under a constitutive promoter. If WT KAP1 was present, the LINE-1 and SVA reporters were both efficiently silenced (**Fig 4C,D**). Mutation of Lys296 in the KAP1 coiled-coil domain, which is involved in multiple electrostatic interactions with the KRAB domain (**Fig 4B**), completely abolished repression of both reporters. Similarly, mutation of Met297 or Leu300, which form part of the hydrophobic core of the KRAB-KAP1 interface, to serine resulted in loss of silencing (**Fig 4C,D**). A third hydrophobic patch mutant, V293S, was not expressed at detectable levels in KAP1 KO HEK 293T cells, but we showed previously that combining all four KAP1 mutations listed above (V293S/K296A/M297A/L300S) abolishes KRAB binding and transcriptional repression (Stoll *et al*., 2019). Together, these data indicate that Leu296, Met297 and Leu300 are required for KRAB binding and repression by KAP1.

### Disruption of the KAP1-KRAB interface affects H3K9 trimethylation genome-wide

Deposition of the repressive epigenetic mark H3K9me3 by SETDB1 is an essential component of targeted KAP1-dependent silencing by KRAB-ZFPs and via CRISPRi (Schultz *et al*., 2002; Thakore *et al*., 2015). To assess the importance of the KRAB binding residues in KAP1 in histone H3K9-trimethylation, we measured the genome-wide distribution of H3K9me3 in cells expressing different KAP1 variants with the CUT&RUN (Cleavage Under Targets and Release Using Nuclease) epigenomic profiling method (Skene *et al*, 2018). We observed a massive loss of H3K9me3 genome-wide in KAP1 KO cells, representing at least 65% of mappable H3K9me3 peaks (**Figs 5A,B and S4**). For example, H3K9me3 was significantly depleted over thousands of retrotransposons from all classes, including full-length primate-specific LINE-1 subfamilies, SVAs and LTR elements (**Fig 5C,D**). The extent of H3K9me3 loss in human cells upon KAP1 depletion is comparable to ChIP-seq data from mouse ESCs (Coluccio *et al*, 2018; Jang *et al*, 2018). Notably, H3K9me3 was more modestly reduced at sites targeted by the HUSH complex, primarily intronic LINE-1s and long exons (Douse *et al*, 2020; Seczynska *et al*, 2022) (**Fig S3**). Complementation of KAP1 KO cells with wild-type KAP1 robustly restored H3K9me3 levels. However, the KRAB-binding mutant CCmut, bearing mutations K296S/M297S/L300S/V293S in the coiled-coil domain, completely failed to restore H3K9 methylation at KAP1-regulated loci (**Fig 5**) despite being expressed in the nucleus at comparable levels to the WT protein (**Fig S5**). Taken together, the data demonstrate that the KRAB-binding residues of KAP1 are required for KAP1-dependent H3K9 methylation.

**Figure 5.**
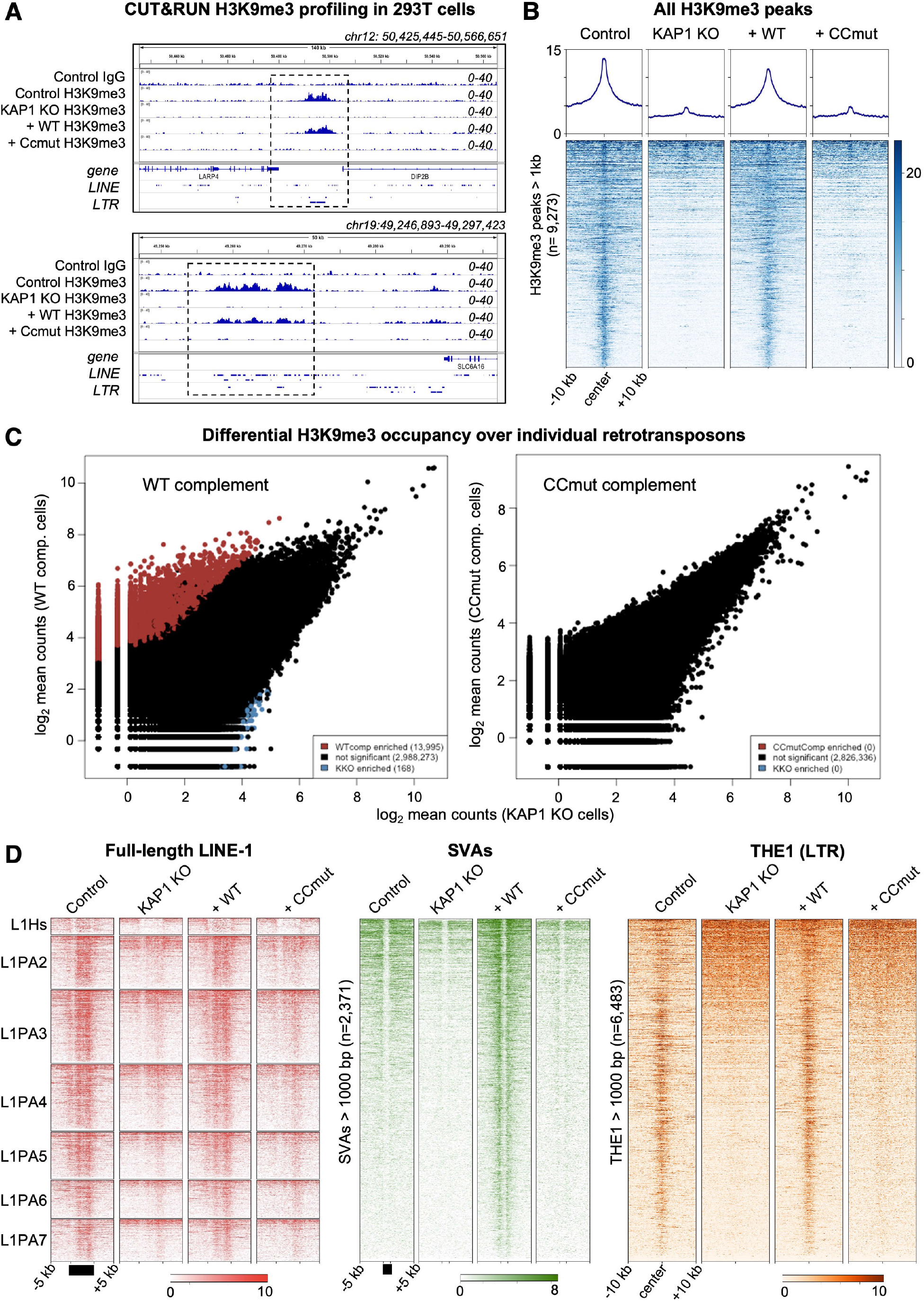
Genome-wide analysis of H3K9me3 distribution in cells expressing wild-type and KRAB-binding-deficient variants of KAP1. See also **Figs S4 and S5**. **(A)** Example genome browser snapshots of H3K9me3 distribution over the hg38 reference in the presence of different KAP1 variants. H3K9me3 distribution is shown at a HERVK9-int LTR element (upper) and a cluster of LINE-1 elements (lower). A control IgG track from parent HEK293T cells is shown for comparison. WT, wild-type; CCmut, KRAB binding-deficient KAP1 variant. Only reads mapping uniquely (MAPQ >10) were retained. Scales are in RPKM, reads per kilobase per million. Experiments were run in duplicate with similar results. **(B)** Heatmaps and summary plots illustrating H3K9me3 levels over H3K9me3 peaks genome wide, in cells expressing different KAP1 variants. Peaks were called on Control cells using SEACR in stringent mode (Meers *et al*, 2019). The summary plots illustrate mean values for each sample. Only reads mapping uniquely (MAPQ >10) were retained. **(C)** Pairwise quantifications of H3K9me3 CUT&RUN counts for KAP1-complemented cells versus KAP1 KO cells over reference retrotransposons (RepeatMasker) generated with DESeq2 (Love *et al*, 2014). Red, H3K9me3 enriched in complemented cells; blue, H3K9me3 enriched in KAP1 KO cells. Enrichment cutoff: p < 0.05; log-fold change ±3. **(D)** Heatmaps showing H3K9me3 CUT&RUN signal enrichment over full-length (>6 kb) LINE-1 subfamilies (left), reference SVAs >1kb (centre) and THE1 LTR elements (right) in cells expressing different KAP1 variants. H3K9me3 is rescued upon complementation with WT KAP1 but not CCmut KAP1. Only reads mapping uniquely (MAPQ >10) were retained; note that only the flanking regions of SVAs are mappable even with a 2 × 150-bp paired end sequencing strategy. Experiments were run in duplicate with similar results.

## Discussion

Here, we report the crystal structure of the core complex between KAP1 and the KRAB domain from a representative KRAB-ZFP, the human LINE-1 repressor ZNF93. KRAB-ZFPs have grown into the largest family of mammalian transcriptional factors, driven by their role in safeguarding the genome from TEs. In parallel, KRAB-ZFPs have acquired vital functions in controlling gene expressing during vertebrate development. A KRAB-dCas9 fusion protein is the key reagent underpinning the CRISPRi technology (Alerasool *et al*., 2020; Gilbert *et al*., 2014; Thakore *et al*., 2015). Our structure provides a detailed and complete three-dimensional atlas of the KRAB-KAP1 binding interface. The functional importance of this interface is validated by our repression reporter assay and epigenomic profiling data.

In the crystal structure, the electron density for the KRAB domain was weaker than for KAP1, potentially indicating some residual degree of conformational flexibility or heterogeneity in the KRAB domain bound to KAP1. Due to the 2:1 stoichiometry of the KAP1:KRAB complex, the KRAB domain could in principle bind to the KAP1 dimer in two equivalent orientations, related by the dimer dyad. However, there is no evidence in the crystal structure of KRAB binding to the KAP1 RBCC dimer in an alternative orientation or conformation (related by the dimer dyad or not). This could be due to steric hindrance from the crystal packing preventing the KRAB from binding in the dyad-related orientation.

Alternatively, the KAP1 dimer may have some inherent asymmetry such that a single binding orientation of the KRAB is favored. Supporting the latter, solution biophysics studies showed that full-length KAP1 forms asymmetric dimers that bind HP1 with a 2:1 KAP1:HP1 stoichiometry (Fonti *et al*, 2019) The HP1 binding domain is located in the C-terminal half of KAP1, outside the RBCC (Schultz *et al*., 2002). Together, the available data support the model that the KAP1 dimer has some degree of intrinsic asymmetry, which is functionally important in that it determines the 2:1 stoichiometry of the complexes with KRAB-ZFPs and HP1.

The KRAB-B box of ZNF93 was present in the crystallized construct but there were no interpretable features in the electron density for the KRAB-B box. AlphaFold2 predicts with medium confidence (pLDDT = 70-90) that the first half of the KRAB-B sequence forms a two-turn helix, which packs against the C-terminal KRAB-A helix, on the opposite side from the KAP1 binding surface (**Fig 1**). A weak electron density peak was present in the KRAB-KAP1 crystal structure (in the *F*_o_ – *F*_c_ Fourier difference map), near where this KRAB-B helix would be expected based on the AlphaFold2 prediction. However, the density was too weak to allow a model to be built, indicating that the KRAB-B box is mobile or disordered in the KRAB-KAP1 complex. Consistent with this, systematic mutagenesis of the KRAB-B box had little effect on KAP1 recruitment, with mutations at only one KRAB-B residue (Pro59) showing a weak effect (Tycko *et al*., 2020).

KRAB-KAP1 complexes recruit multiple effectors that modify the and epigenetic conformational landscape of chromatin target loci including HP1, SETDB1, NuRD and SMARCAD1. Except for SMARCAD1, these effectors bind to sites outside the KAP1 RBCC, in the C-terminal half of KAP1. SMARCAD1 is an ATP-dependent chromatin remodeler thought to be important to generate and maintain the necessary chromatin conformation in pluripotent embryonic stem cells (Ding et al., 2018). The KRAB-KAP1 crystal structure shows that the KRAB and SMARCAD1 CUE1 domains bind KAP1 RBCC dimers independently, without steric interference (**Fig 2**). Hence the KRAB binding site on KAP1 is distal from all known effector binding sites on KAP1, meaning that all KAP1 effectors are expected to be recruited to KRAB-ZFP chromatin binding sites.

By recruiting chromatin modifying proteins to KRAB-ZFP binding sites, KAP1 protects the genome from invasion by TEs and plays a key role in regulating gene expression in early development, across tetrapod vertebrate species. The KRAB-KAP1 interaction lies at the nexus of this vital transcriptional control axis. Our work identifies and functionally validates the KRAB-KAP1 molecular interface. The KRAB-KAP1 structure will guide future efforts to further increase the repression potency of KRAB domains in CRISPRi and potentially other applications. The ability to understand and manipulate how, and when KAP1 is recruited to its target loci by KRAB-ZFPs, specifically in stem cells or pluripotent cells, could create new opportunities to direct cell fate determination, for example in cell therapy applications.

## Materials and Methods

### Cell lines and bacterial strains

HEK293T cells were a kind gift from Helen Rowe (University College London). Selenomethionine-labelled proteins were expressed in *Escherichia coli* B834(DE3) cells (Merck). All other proteins were expressed in *Escherichia coli* BL21(DE3) cells (New England BioLabs).

### Expression vectors

A synthetic gene encoding SMARCAD1 CUE1 (residues 151-198; UniProt Q9H4L7) codon-optimized for *Escherichia coli* (*E. coli*) was cloned into the first multiple cloning site (MCS) of the pRSFDuet plasmid (Novagen), with N-terminal hexahistidine purification (His_6_) tag followed by a TEV protease cleavage site. The T4L-RBCC fusion construct was described previously (Stoll *et al*., 2019). To improve protein crystallization the B-box 1 domain (residues 141-202 of KAP1) was deleted, generating the T4L-RBCCβB1 plasmid. A codon-optimised gene encoding residues 2-71 of ZNF93 (UniProt P35789) was cloned into MCS1 of pCDFDuet, preceded by Twin-StrepII and MBP purification tags and a HRV 3C protease cleavage site.

For expression in mammalian cells, full-length KAP1 (UniProt Q13263) with triple FLAG tag was cloned into pLEXm as described previously (Stoll *et al*., 2019). For lentivirus production, KAP1 was subcloned into pHRSIN.

### Protein expression and purification

T4L-RBCCβB1-ZNF93 KRAB complexes were produced by co-expression in *E. coli* BL21 (DE3) cells (New England BioLabs). Cells were grown at 37°C in 2×TY medium. When the optical density (OD_600_) of the cultures reached 0.4-0.5, the culture medium was supplemented with 50 µM ZnSO_4_ and the incubator temperature was lowered to 16°C. At an OD_600_ of 0.8, protein expression was induced with 0.2 mM isopropyl-β-D-thiogalactopyranoside (IPTG) for 18 h, before the cells were harvested by centrifugation (6,000×g for 15 min). The bacteria pellets were resuspended in wash buffer (50 mM Tris pH 8, 0.2 M NaCl, 0.5 mM TCEP) supplemented with 1:10,000 (v/v) Benzonase nuclease (Sigma) and 1×cOmplete EDTA-free protease inhibitors (Roche) and lysed by sonication. The lysate was clarified by centrifugation (30 min, 40,000×g) and applied to a 5-ml StrepTrap HP column (Cytiva) equilibrated in wash buffer. The column was washed with 30 column volumes (CV) of wash buffer before bound proteins were eluted with wash buffer supplemented with 3 mM D-desthiobiotin (Sigma). The protein was transferred into wash buffer without D-desthiobiotin and incubated overnight at 4°C with 1:100 (w/w) HRV 3C protease to remove the Twin-StrepII-MPB tag. Uncleaved protein and free tag were subsequently captured using a StrepTrap HP column and the sample was further purified by size-exclusion chromatography using a HiLoad (16/600) Superdex 200 pg column (Cytiva) equilibrated in 20 mM HEPES pH 8, 0.5 M NaCl, 0.5 mM TCEP. Selenomethionine labelled proteins were expressed in *E. coli* B834 (DE3) cells (Novagen) and purified like the native protein, except that all buffers contained 1 mM TCEP.

*E. coli* BL21 (DE3) cells were transformed with the His_6_-CUE1 pRSFDuet plasmid and grown at 37°C. At OD_600_ = 0.8, the incubator temperature was reduced to 18°C and protein expression was induced with 0.2 mM IPTG for 18 h. The cells were harvested by centrifugation, resuspended in wash buffer (50 mM Tris pH 8, 0.3 M NaCl, 20 mM imidazole, 0.5 mM TCEP) supplemented with 1:10,000 (v/v) Benzonase nuclease (Sigma) and 1×cOmplete EDTA-free protease inhibitors (Roche) and lysed by sonication. Insoluble cell debris was removed by centrifugation (30 min, 40,000×g) and the supernatant was applied to a 5-ml HisTrap HP column (Cytiva) equilibrated in wash buffer. The column was washed with 30 CV of wash buffer before bound proteins were eluted with elution buffer (50 mM Tris pH 8, 0.3 M NaCl, 250 mM imidazole, 0.5 mM TCEP). The eluted protein was further purified by size-exclusion chromatography using a HiLoad (26/600) Superdex 75 pg column (Cytiva) equilibrated in 20 mM HEPES pH 8, 0.2 M NaCl.

### X-ray crystallography

Crystals were grown at 18°C by sitting drop vapor diffusion. Purified T4L-RBCCΔB1 - ZNF93 KRAB complex (5 g L^-1^, 43 µM) was mixed with SMARCAD1 CUE1 domain (5 g L^-1^, 610 µM) at a 1:2.2 molar ratio. The sample was subsequently mixed with an equal volume of reservoir solution optimized from the Index screen (Hampton Research): 11% (w/v) PEG 5000 MME, 5% Tacsimate, 0.1 M HEPES pH 7. Crystals appeared after 2 days and were frozen in liquid nitrogen with 20% glycerol as a cryoprotectant. X-ray diffraction data were collected at 100 K at Diamond Light Source (beamline i04) and processed with xia2 (DIALS, AIMLESS). The structure was solved by molecular replacement using T4L-fused KAP1 RBCC (PDB 6QAJ) (Stoll *et al*., 2019) as a search model. The atomic models for the SMARCAD1 CUE1 domain (PDB 6QU1) (Lim *et al*., 2019) were docked into the phased electron density map. The ZNF93 KRAB domain was built using the Alphafold2 prediction of ZNF93 and the NMR structure of a mouse KRAB domain (PDB 1V65) (Saito *et al*., 2003) as guides. The model was iteratively refined using COOT and PHENIX (Adams *et al*., 2011). See **Table 1** for data collection and refinement statistics.

Selenomethionine-labelled point mutants crystallized in similar conditions as the native WT complex and were cryoprotected with either 20% glycerol or 25% ethylene glycol prior to freezing in liquid nitrogen. Data was collected at a wavelength of 0.9795 Å. The structures were solved by molecular replacement using the structure of the T4L-RBCCΔB1 - ZNF93 KRAB - CUE1 complex. The models were refined using the LORESTR pipeline (Nicholls *et al*, 2017) implemented in CCP4 (Cowtan *et al*, 2011) and PHENIX. Selenium sites were located using Phaser MR-SAD in PHENIX (McCoy *et al*, 2007).

### Transcriptional silencing assay

The transcriptional silencing activity of KAP1 mutants was measured using reporter plasmids in which SVA or LINE-1 sequence upstream of a minimal SV40 promoter enhances firefly luciferase activity unless KAP1 and the cognate KRAB-ZFP (ZNF91 and ZNF93, respectively) are present to repress the reporter. KAP1 KO HEK293T cells in 24-well plates were cotransfected with 20 ng firefly luciferase reporter plasmid, 0.2 µg plasmid encoding ZNF91 or ZNF93, 0.2 µg pLEXm plasmid encoding WT or mutant KAP1, and 0.4 ng plasmid encoding Renilla luciferase using FuGENE 6 (Promega). Luciferase activity was measured 48 h post-transfection using the Dual Luciferase assay kit (Promega) and a PHERAstar FSX microplate reader (BMG Labtech). Replicates were performed on separate days. Firefly luciferase values were normalized to *Renilla* luciferase values to control for transfection efficiency. Statistical significance was assessed with an unpaired t test (assuming Gaussian distributions, without Welch’s correction) with PRISM 9 (GraphPad).

### Western Blotting

4 × 10^5^ HEK293T cells were lysed in 100 μl of Passive Lysis Buffer (Promega). 10 μl of cell lysates were separated on a NuPAGE 4-12% Bis-Tris polyacrylamide gel (ThermoFisher). The samples were transferred onto nitrocellulose membranes using the iBlot2 Dry Blotting System (ThermoFisher). The membrane was blocked with 5% (w/v) skim milk powder (Sigma-Aldrich) in PBS for 1 h at room temperature before it was incubated overnight at 4°C with primary antibody diluted in PBS-T (PBS with 0.1% Tween-20) containing 5% (w/v) skim milk powder. Rabbit anti-KAP1 antibody (Abcam, cat. no. ab10484, RRID:AB_297223) was diluted 1:10,000; rabbit anti-actin antibody (Abcam, cat. no. ab219733; RRID:AB_219733) was diluted 1:2,000. Subsequently, the membrane was washed four times with PBS-T and incubated with DyLight 800 goat anti-rabbit IgG (Cell Signaling Technology, cat. no. 5151, RRID:AB_10697505) diluted 1:10,000 in PBS-T containing 5% (w/v) skim milk powder. After 30 min at room temperature, the membrane was washed four times with PBS-T, twice with PBS and once with ultrapure water. Blots were imaged using an Odyssey CLx gel scanner (LI-COR Biosciences).

### Generation of stable cell lines

Lentivirus was produced by transfecting HEK23T cells in 6-well plates with 0.6 µg pMD2.G, 1.2 µg p8.91 and 1.2 µg KAP1 pHRSIN using FuGENE 6 (Promega). Virus-containing supernatant was collected 48 h post-transfection, filtered through a 0.45 µm PVDF membrane and used to infect KAP1 KO HEK239T cells. Transduced cells were selected with 200 µg ml^-1^ hygromycin.

### Surface plasmon resonance (SPR)

SPR was performed on a Biacore T200 system with Series S CM5 sensor chips (Cytiva). Reference and sample channels were equilibrated in 20 mM HEPES pH 8.0, 0.5 M NaCl, 0.5 mM TCEP at 20°C. MBP-KRAB was immobilized onto the sensor chip until a response unit (RU) value of approximately 300 (ZNF93 WT) or 600 (ZNF93 E35R) was reached. Analytes in 1:2 dilution series at an initial concentration of 40 µM were injected for 120 s followed by a 900 s dissociation phase. After each injection cycle, the sensor surface was regenerated with 20 mM NaOH for 30 s with a 120-s post-regeneration stabilization period. Data were fitted using a biphasic kinetic model with PRISM 9 (GraphPad) to determine *k*_on_, *k*_off_ and *K*_d_.

### CUT&RUN H3K9me3 profiling

We followed the protocol detailed by Henikoff and colleagues (Skene *et al*., 2018). Briefly, 250,000 cells (per antibody/cell line combination) were washed twice (20 mM HEPES pH 7.5, 0.15 M NaCl, 0.5 mM spermidine, 1x Roche complete protease inhibitors) and attached to ConA-coated magnetic beads (Bangs Laboratories) pre-activated in binding buffer (20 mM HEPES pH 7.9, 10 mM KCl, 1 mM CaCl_2_, 1 mM MnCl_2_). Cells bound to the beads were resuspended in 50 µl buffer (20 mM HEPES pH 7.5, 0.15 M NaCl, 0.5 mM Spermidine, 1x Roche complete protease inhibitors, 0.02% w/v digitonin, 2 mM EDTA) containing primary antibody (1:100 dilution). Incubation proceeded at 4°C overnight with gentle shaking. Tubes were placed on a magnet stand to remove unbound antibody and washed three times with 1 ml digitonin buffer (20 mM HEPES pH 7.5, 0.15 M NaCl, 0.5 mM Spermidine, 1x Roche complete protease inhibitors, 0.02% digitonin). pA-MNase (35 ng per tube, a generous gift from Steve Henikoff) was added in 50 µl digitonin buffer and incubated with the bead-bound cells at 4°C for 1 h. Beads were washed twice, resuspended in 100 µl digitonin buffer and chilled on ice. Genome cleavage was stimulated by addition of 2 mM CaCl_2_, briefly vortexed and incubated on ice for 30 min. The reaction was quenched by addition of 100 µl 2x stop buffer (0.35 M NaCl, 20 mM EDTA, 4 mM EGTA, 0.02% digitonin, 50 ng/µl glycogen, 50 ng/µl RNase A, 10 fg/µl yeast spike-in DNA (a generous gift from Steve Henikoff)) and vortexing. After 10 min incubation at 37°C to release genomic fragments, cells and beads were pelleted by centrifugation (16,000 g, 5 min, 4°C) and fragments from the supernatant purified with a Nucleospin PCR clean-up kit (Macherey-Nagel). Illumina sequencing libraries were prepared using the Hyperprep kit (KAPA) with unique dual-indexed adapters (KAPA), pooled and sequenced on a NovaSeq6000 instrument. Paired-end reads (2×150) were aligned to the human genome (hg38) using Bowtie2 (--local –very-sensitive-local –no-mixed –no-discordant –phred33 -I 10 -X 700) and converted to bam files with samtools. Only reads with unique mapping (MAPQ>10) were retained for all analyses. PCR duplicates were removed with Picard. Peaks were called on control cells using SEACR in stringent mode (Meers *et al*., 2019). Differential H3K9me3 occupancy was assessed qualitatively with deepTools (computeMatrix and plotHeatmap commands on RPKM-normalized bigwig coverage files) and quantitatively with a combination of featureCounts, HOMER (Heinz *et al*, 2010) and DESeq2 (Love *et al*., 2014). Retrotransposon annotations were downloaded from the RepeatMasker hg38 database. CUT&RUN experiments were done in two independent replicate experiments.

### Immunofluorescence microscopy

Cells were grown in 24-well plates on poly-L-lysine coated coverslips, fixed with 4% formaldehyde in PBS for 15 min and permeabilized with 0.1% Triton X-100 in PBS. After blocking with PBS and 10% FBS for 1 h, samples were incubated for 1 h with the primary antibody (mouse anti-KAP1, Proteintech, cat. no. 66630-1-Ig, RRID:AB_2732886) diluted 1:500 in PBS and 10% FBS. The cells were washed three times with PBS and 10% FBS and then incubated for 1 h with the secondary antibody (Alexa Fluor 488 goat anti-mouse, ThermoFisher, cat. no. A-11001, RRID:AB_2534069) diluted 1:500 in PBS and 10% FBS. After three washes with PBS and 10% FBS, one wash with PBS, and one wash with water, the coverslips were mounted with ProLong Gold Antifade Mountant with DAPI (ThermoFisher). Images were acquired using a Zeiss LSM 780 confocal microscope with a 63x/1.4 NA oil immersion objective.

### Quantification and statistical analysis

No statistical methods were used to predetermine sample size, experiments were not randomized, and the investigators were not blinded to experimental outcomes. Luciferase-reporter cell signaling data are represented as the mean ± standard error of the mean of three replicates conducted in a single independent experiment. Data are representative of at least three independent experiments.

## Data availability

The atomic coordinates were deposited in the Protein Data Bank with code PDB:7Z36. The original experimental X-ray diffraction images were deposited in the SBGrid Data Bank (data.SBGrid.org), with Data ID 880, DOI:10.15785/SBGRID/880. Other data are available from the corresponding author upon reasonable request. The CUT&RUN data were deposited in the Gene Expression Omnibus (GEO) database under accession code GSEXXXXX.

## Acknowledgments

KAP1 KO cells and plasmids for the transcriptional silencing assay were kind gifts from Helen Rowe (University College London). The plasmids were provided with permission from David Haussler (Univ. of California Santa Cruz). pA-MNase used in CUT&RUN experiments was a kind gift from Steven Henikoff (Fred Hutchinson Cancer Research Center). Crystallographic data were collected on beamline i04 at Diamond Light Source (DLS). Access to DLS (proposal MX21426) was supported by the Wellcome Trust, MRC and BBSRC. This work was supported by MRC research grant MR/S021604/1 to Y.M., Wellcome Trust Senior Research Fellowship 217191/Z/19/Z to Y.M., Wellcome Trust PhD Studentship 205833/Z/16/Z to G.S., and Swedish Society for Medical Research (Svenska Sällskapet för Medicinsk Forskning) grant S19-0100 to C.H.D.

## Author contributions

G.A.S. and Y.M. conceived the study. G.A.S. purified the proteins and performed the cell reporter assays, biochemical assays, and crystallographic structure determination. N.P. and C.H.D. performed and analysed the CUT&RUN experiments. G.A.S. and Y.M. wrote the manuscript with discussion and input from N.P. and C.H.D.

## Conflict of interest

Y.M. is a consultant for Related Sciences LLC.

## Supplementary Figures and Figure Legends

**Supplementary Figure 1.**
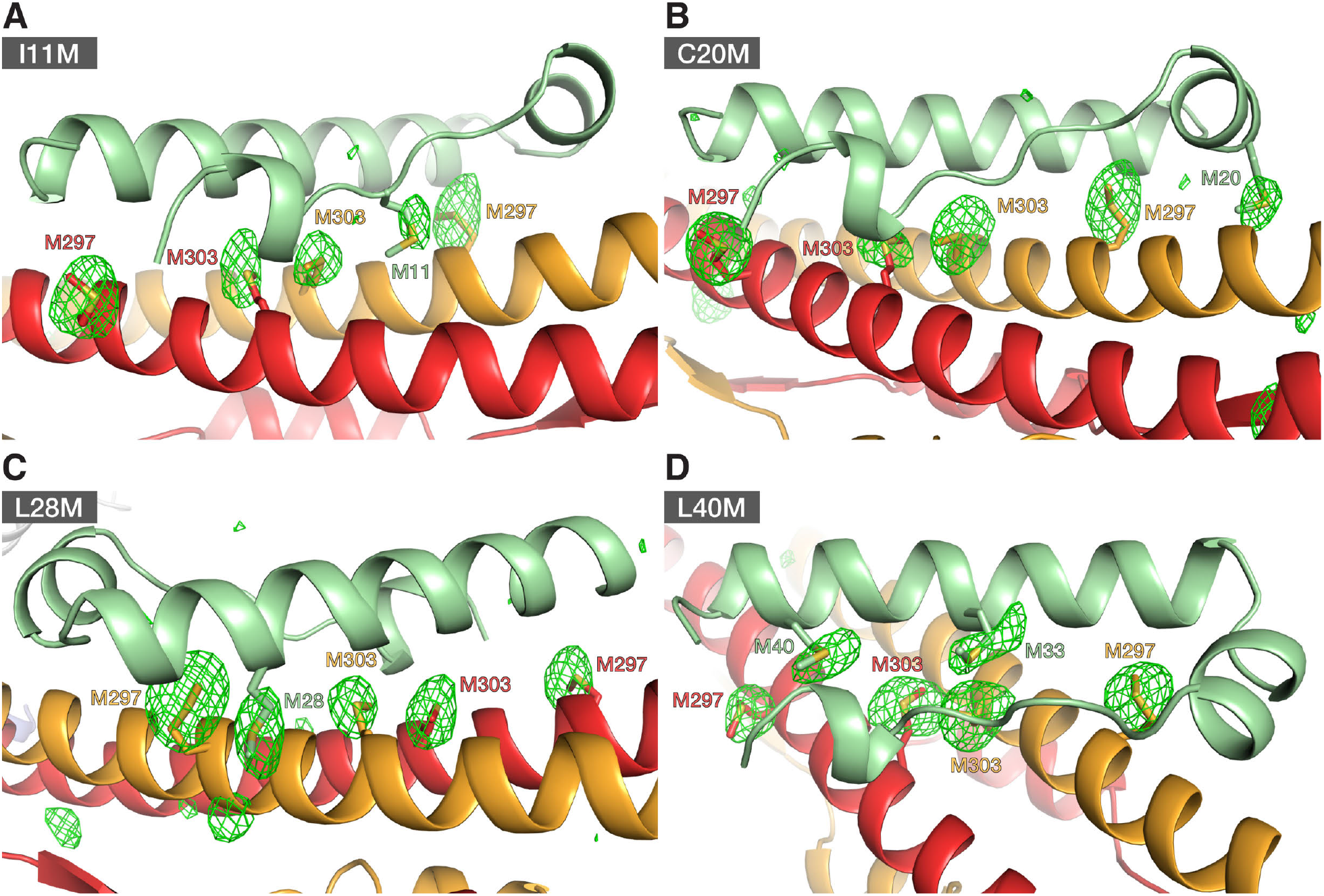
Anomalous Fourier maps of selenomethionine derivatives of KAP1 RBCC in complex with four different ZNF93 methionine-insertion mutants, Related to **Figure 1**. (**A-D**) X-ray diffraction data were collected at the selenium K absorption edge. Anomalous Fourier maps were contoured at 3.5 σ. The Fourier maps show the positions of selenium atoms of the selenium-substituted methionine residues the KRAB and KAP1 RBCC domains. **(A)** KAP1 RBCC in complex with the ZNF93 KRAB I11M mutant. **(B)** KAP1 RBCC in complex with the ZNF93 KRAB C20M mutant. **(C)** KAP1 RBCC in complex with the ZNF93 KRAB L28M mutant. **(D)** KAP1 RBCC in complex with the ZNF93 KRAB L40M mutant.

**Supplementary Figure 2.**
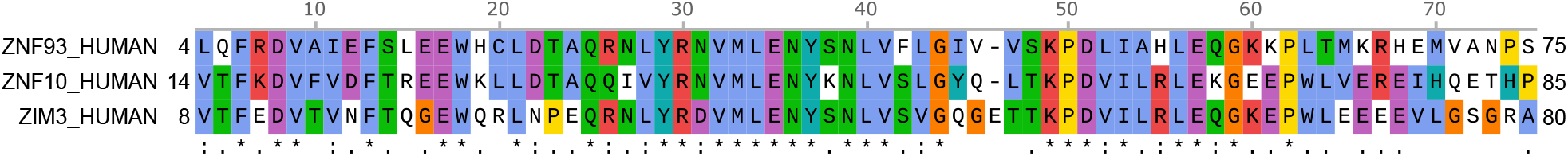
Amino acid sequence alignment of the KRAB domains from ZNF93, ZNF10 and ZIM3, Related to **Figure 3**. Fused to catalytically inactive Cas9, the KRAB domains of ZNF10 and ZIM3 are used for potent gene repression that can be programmed in a sequence-specific manner via the CRISPRi approach (Alerasool et al., 2020; Gilbert et al., 2014; Thakore et al., 2015).

**Supplementary Figure 3.**
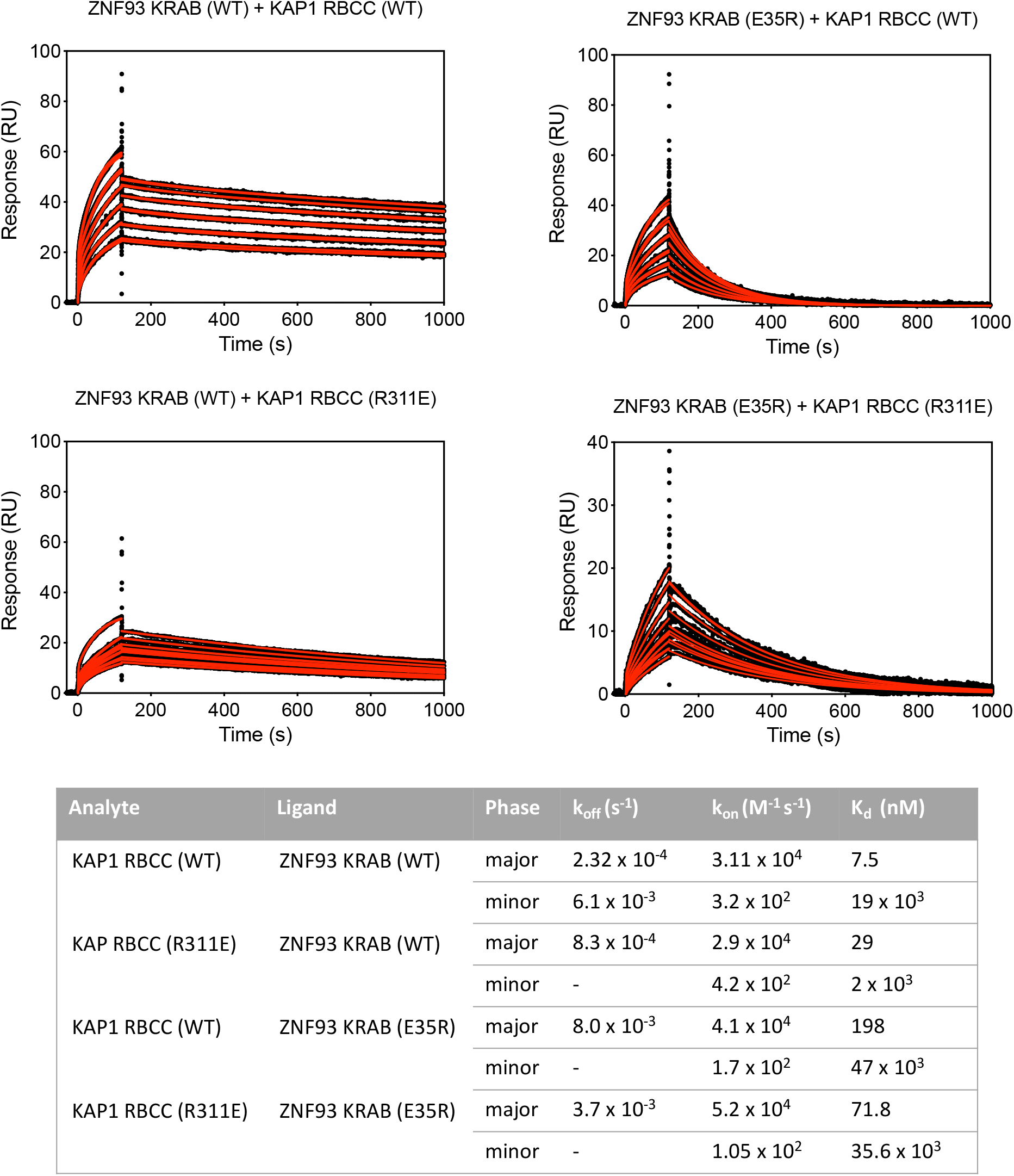
Surface plasmon resonance (SPR) KAP1-KRAB binding assay titration curves, Related to **Figure 4**. Upper panel: SPR sensorgrams for KAP1 binding to immobilized MBP KRAB. The fits for the association and dissociation kinetics are shown in red. Lower panel: Data were fitted using a biphasic kinetic model with PRISM 9 (GraphPad) to determine rate constants (*k*_on_, *k*_off_) and binding affinities (*K*_d_).

**Supplementary Figure 4.**
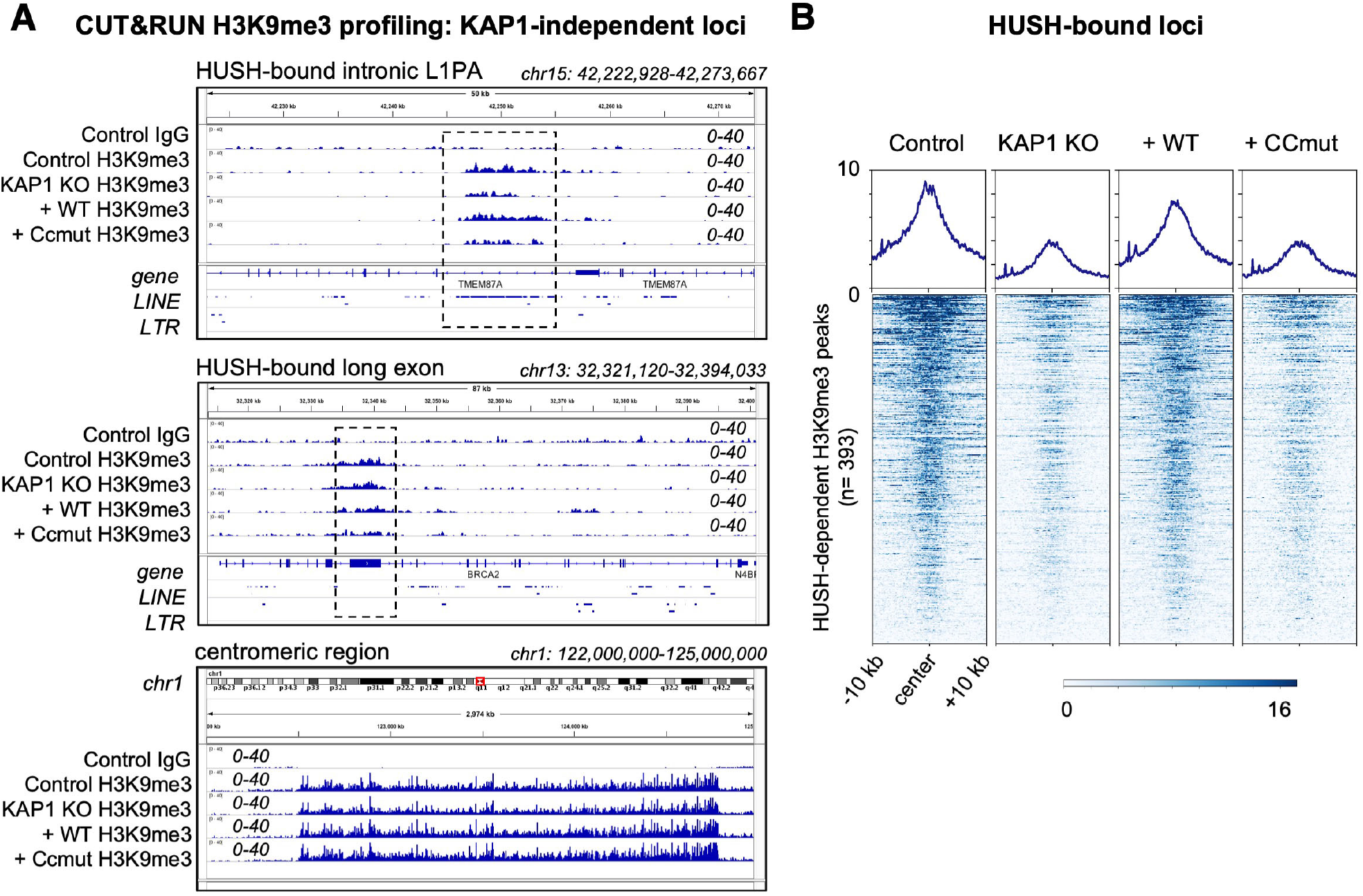
Genome-wide analysis of H3K9me3 distribution in cells expressing wild-type and KRAB-binding-deficient variants of KAP1, Related to **Figure 5**. **(A)** Representative H3K9me3 distribution (over the hg38 reference) in the presence of different KAP1 variants at three different types of KAP1-independent loci: an intronic LINE-1 (L1PA) element bound by the HUSH complex (upper); a HUSH-bound long exon (middle); and a centromeric region (lower). A control IgG track from parent HEK293T cells is shown for comparison. WT, wild-type; CCmut, KRAB binding-deficient KAP1 variant. Only reads mapping uniquely (MAPQ >10) were retained. Scales are in RPKM, reads per kilobase per million. Experiments were run in duplicate with similar results. **(B)** Heatmaps and summary plots illustrating H3K9me3 levels over H3K9me3 peaks at HUSH-bound loci (Douse et al., 2020; Seczynska et al., 2022), in cells expressing different KAP1 variants. Peaks were called on Control cells using SEACR in stringent mode (Meers et al., 2019). The summary plots illustrate mean values for each sample. Only reads mapping uniquely (MAPQ >10) were retained.

**Supplementary Figure 5.**
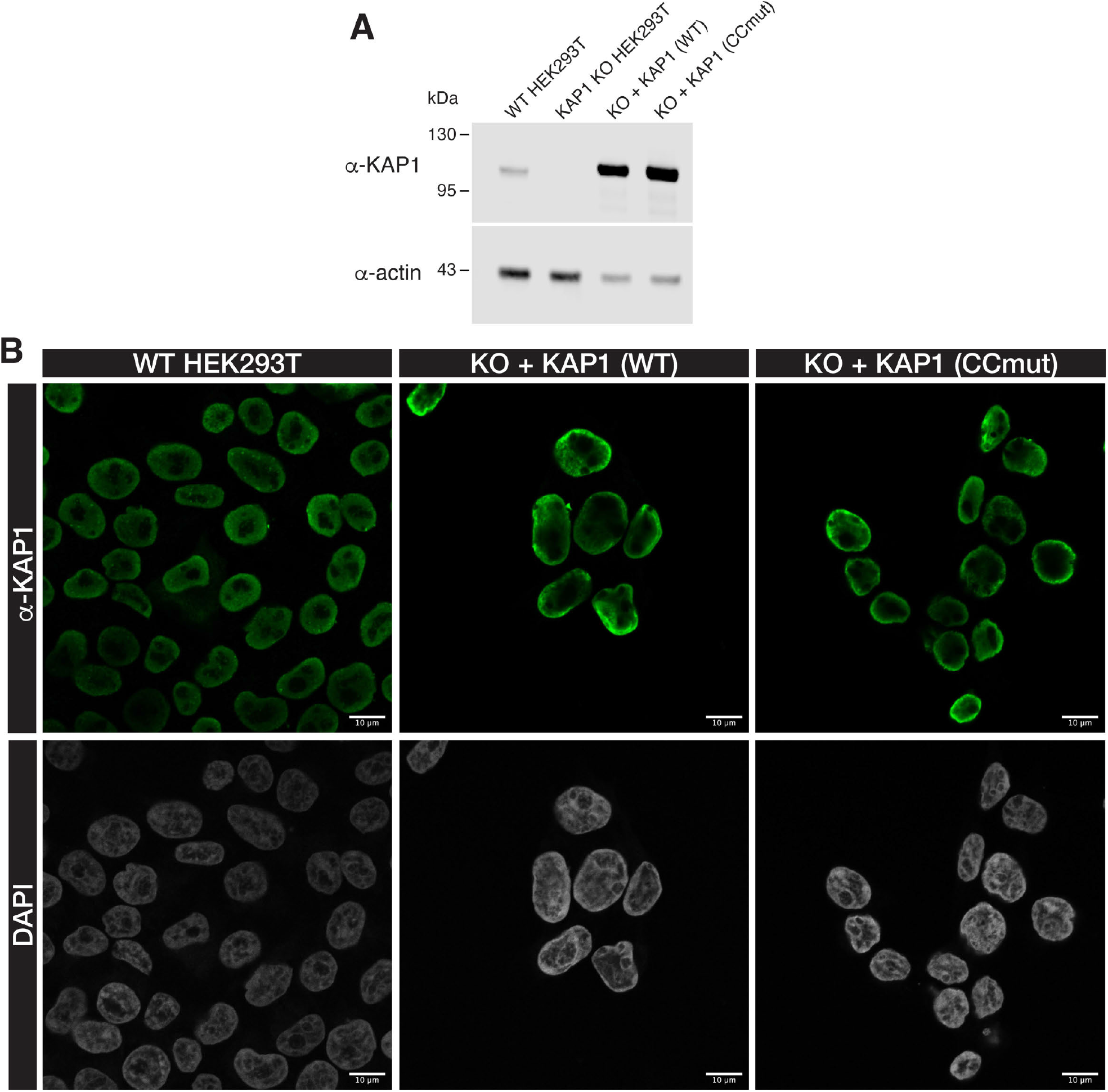
Expression levels of KAP1 variants in HEK239T cells, Related to **Figure 5**. **(A)** Western blot of the HEK293T cells used for CUT&RUN genomic profiling. WT, wild-type; KO, KAP1 knockout; CCmut, KRAB binding-deficient KAP1 variant K296S/M297S/L300S/V293S. **(B)** Confocal immunofluorescence microscopy of HEK293T cells used for CUT&RUN genomic profiling stained with anti-KAP1 antibody and DAPI nuclear stain. Scale bars, 10 µm.

